# Single-Molecule Proteomics via a Dynamic Translocase and Physics-Informed Machine Learning

**DOI:** 10.64898/2026.08.17.745284

**Authors:** Jaylen E. Taylor, Parichit Sharma, Bryan A. Krantz

**Author notes:** **Email** (BAK).

## Abstract

Single-molecule protein sequencing promises to democratize clinical proteomics, but platforms retrofitting static DNA-sequencing nanopores face a fundamental biophysical bottleneck: they only measure one-dimensional excluded volume. Consequently, these static calipers struggle to resolve isobaric residues, requiring complex DNA-handle chemistries and target concentrations that exceed clinically relevant abundance ranges. Here, we introduce a dynamical, target-docking translocase engine – the anthrax toxin protective antigen (PA) – as a label-free single-molecule peptide sensor. By extracting the multi-state thermodynamic friction generated as the pore’s active site dynamically “breathes” around translocating analytes, we trained a physics-informed machine learning (PIML) architecture to classify a 20-member guest-host peptide library panel representing all 20 canonical amino acids at the single-event level. Operating at low nanomolar concentrations under a 35-millisecond thermodynamic read constraint, the translocase resolved isobaric variants (leucine and isoleucine). Furthermore, we achieved 98.02 (±0.05)% classification accuracy on a panel of five un-tagged, native clinical biomarkers (e.g., KRAS G12D, angiotensin, bradykinin). Transitioning from static volumetric measurement to time-domain thermodynamic fingerprinting establishes the requisite protein nanopore hardware for *de novo* proteomics.

## Introduction

The democratization of genomics was driven by the advent of single-molecule nanopore sequencing, a technology optimized around the translocation of uniformly charged nucleic acids through rigid, static biological pores. However, the translation of this architecture to single-molecule proteomics has stalled against a fundamental biophysical bottleneck. The current field relies heavily on a “sunk cost” infrastructure, attempting to retrofit static nanopores (e.g., α-hemolysin (*1, 2*), aerolysin (*3*), MspA (*4*), and CsgG (*5, 6*)) to read structurally folded, chemically diverse polypeptides. This approach to single-molecule proteomics is biophysically flawed. A static nanopore acts as a passive spatial caliper, reducing the measurement of a complex proteome to a single dimension: excluded volume. Consequently, passive pores mathematically struggle to resolve isosteric variants, post-translational modifications (PTMs), and isobaric residues (e.g., leucine and isoleucine) that produce identical volumetric footprints.

To overcome this one-dimensional limit, we propose a paradigm shift toward dynamical nanopore translocases. Rather than a passive conduit, the anthrax toxin protective antigen (PA) forms a highly evolved, ∼0.5 MDa homooligomeric (*7-9*) protein-handling, ratcheting nanomachine (*10*) and model system (*11*). The PA translocase is a self-contained engine equipped with multiple active-site peptide clamps (*12, 13*) **(Fig. 1A)**. The α-clamp actively recruits and grips target sequences nonspecifically with extreme sensitivity (*14*), thereby driving the empirical Limit of Detection (LOD) for peptide analytes down into the low nanomolar range. The analyte is subsequently ratcheted through the ϕ-clamp – a radially symmetric ring of phenylalanine residues (F427) serving as the primary active site (*15*) **(Fig. 1B)**. Unlike static pores, the ϕ-clamp acts as a dynamic reading head (*16, 17*). It physically clamps and dilates (*16-18*) around the translocating peptide, “breathing” in response to the specific steric and chemical friction of the analyte.

**Figure 1.**
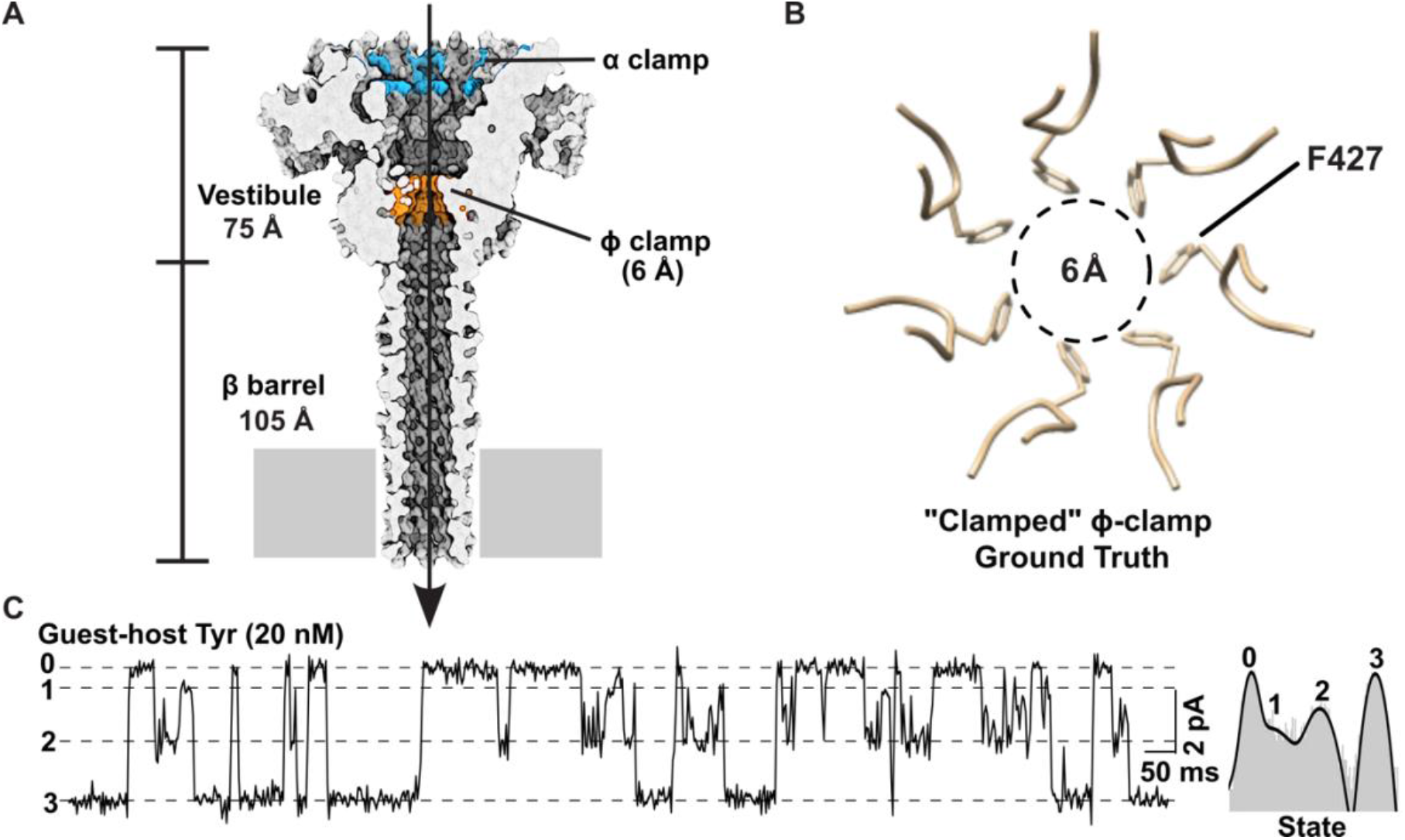
Dynamical PA Nanopore Peptide Biosensor. **(A)** Sagittal view of PA nanopore from atomic-resolution Cryo-EM (*12, 13*). The 105 Å β-barrel and 75 Å vestibule are denoted alongside key functional clamp sites (α-clamp, ϕ-clamp) **(B)** Inset highlighting the dynamic ϕ-clamp (F427) active site (*15*). The resting aperture in the cryo-EM structure exhibits a luminal diameter of ∼6 Å, mathematically aligning with the van der Waals radius of a single residue. **(C)** Representative current signal of PA translocation of guest-host Tyr: cis-added 20 nM guest-host Tyr peptide translocation events (sequence KKKKKYYSYY where guest = Y) at +70 mV (cis positive) in 100 mM KCl, pH 5.6. Discrete conductance states are enumerated: fully blocked (State 0), partially ∼80-90% blocked intermediate 1 (State 1), partially ∼50% blocked intermediate 2 (State 2), and fully open (State 3). Logscale histogram (right) is fitted to four Gaussians. Consistent and reproducible data samples have been reported in Ghosal et al. (*16*), Colby and Krantz (*21*) and results described herein. Sample recordings of the other 20 natural amino acids in the guest-host system are in **Supplemental Figure 1** with corresponding histograms in **Supplemental Figure 2**.

This active engagement between the ϕ-clamp and peptide analyte eliminates the need for the elaborate sample preparation required by static legacy platforms. Because static pores lack an intrinsic active-capture and ratcheting mechanism, the field has been forced to adopt DNA-handle chemistries, covalently ligating massive poly-anionic DNA tails to peptides to orient them or relying on complex, DNA helicase motors to forcibly drag them through the constriction (*3, 19*). These requirements for single-molecule proteomics are clinically disadvantageous and severely limit high-throughput sample processing. By contrast, the PA translocase intrinsically captures and ratchets completely label-free analytes, requiring zero motor accessories or synthetic handles.

Furthermore, the dynamical architecture of PA solves the critical hardware bandwidth imperative. Because static pores rely on minute fluctuations in amplitude to separate analytes, they require extreme sampling frequencies (e.g., 250 kHz to 1 MHz) to suppress thermal noise (*3*). Such high data density is incompatible with the parallelization required for commercial clinical proteomics. Since the PA translocase shifts the discriminative work to the biological machine generating macroscopic conformational state transitions – the necessary telemetry manifests cleanly at low bandwidths (400 to 600 Hz) (*17, 20-22*). This low-frequency requirement ensures the biological sensor is perfectly and natively compatible with the data sampling limits of commercial, high-throughput CMOS-based, nanopore-interfaced sensor arrays (e.g., the Axbio AXP-100 (*23*), Oxford Nanopore MinION (*24*)), as demonstrated in nucleic acid sequencing, rendering the platform highly translational to the clinic.

The continuous clamping and unclamping bistable fluctuations of the ϕ-clamp are not stochastic noise; they are the deterministic output of a thermodynamic energy landscape (*17*). By treating the pore as an active machine, we can extract high-dimensional biophysical features from single peptide translocation events – including Markovian transition matrices, discrete state probabilities, and millisecond dwell times. By pairing these >60-dimensional kinetic fingerprints with physics-informed machine learning (PIML) (*20, 21, 25, 26*), we can mathematically decode the dynamical signal, bypassing the volumetric limits of the static caliper.

The current frontiers of single-molecule proteomics demand the ability to decode the entire “periodic table” of the twenty canonical amino acids, identify clinical biomarkers, resolve more than three hundred PTMs, and ultimately perform *de novo* sequencing of native peptides. Here, we demonstrate that a PIML architecture can decode the twenty natural amino acids in continuous guest-host peptides by utilizing the quasi-thermodynamics and kinetics of their complex multi-state transition matrices with 95.81 (±0.04)% accuracy. Moreover, we extend this framework to the label-free detection of clinically relevant, unmodified disease biomarkers, successfully resolving targets with 1-2 amino acid differences (e.g., KRAS G12D neoantigens, angiotensin metabolites, and bradykinin) with 98.02 (±0.05)% accuracy. Finally, this dynamical progress with the PA nanopore is objectively benchmarked against a legacy static nanopore (*3*) utilizing an uncurated, quantitative machine-learning pipeline to establish the absolute physical limits of one-dimensional volumetric measurement.

To ensure absolute transparency and reproducibility, all source code, machine-learning architectures, and raw data utilized in this study have been made freely available.

## Results

### High-Dimensional Thermodynamic Fingerprinting of the 20 Canonical Amino Acids

To establish a foundational biophysical baseline for the dynamical translocase, we systematically mapped the thermodynamic and kinetic signatures generated by translocation of a guest-host peptide library representing all 20 canonical amino acids. Utilizing high-resolution single-channel planar lipid bilayer electrophysiology, we recorded the translocation of a standardized 10-residue guest-host peptide library (KKKKKXXSXX, where X represents the target amino acid) driven through the wild-type anthrax toxin PA nanopore at a constant +70 mV electrophoretic potential (cis positive) at 20 nM peptide concentration. (Note well that one stereochemical variant, guest-host TrpDL, was included as a 21^st^ peptide, where every other residue of the sequence (i.e., KkKkKwWsWw) was synthesized with d stereochemistry (lower case letters). Unlike static pores that yield one-dimensional amplitude blockades, the PA translocase dynamically “breathes” around the translocating analyte. The physical interaction between the target residue and the 6-Å F427 ϕ-clamp generated rich, multi-state Markovian transitions (fluctuating between fully blocked (State 0), deeply constricted (State 1), dilated (State 2), and open states (State 3)) **(Fig. 1C, Supplementary Figs. 1 and 2)**.

By extracting the millisecond dwell times and state observation frequencies across >600,000 continuous single-molecule events, we mapped a dual-layered energetic fingerprint for each target. First, we computed the quasi-thermodynamic state occupancies, revealing the unique energetic minima dictated by the peptide residing in distinct conformational sub-states **(Fig. 2A, Supplementary Dataset 1)**. Second, we extracted the specific kinetic activation energies (*RT* ln τ_mean_) required to transition between these states **(Fig. 2B, Supplementary Dataset 2)**. As visualized in these high-dimensional heat maps, each amino acid imposes a specific steric, electrostatic, and hydrophobic friction upon the ϕ-clamp. Bulky aromatics (e.g., tryptophan, tyrosine) not only heavily populate the deep blockade states but also face massive kinetic barriers driving abortive cycling (State 0 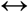 State 2 transitions). Conversely, small aliphatic residues (e.g., alanine) slide through the vestibule with minimal thermodynamic resistance and low transition barriers. This confirms that the dynamical translocase functions not as a passive volumetric caliper, but as a highly sensitive, time-domain thermodynamic sensor – extracting both unique state populations and discrete transition kinetics for every amino acid.

**Figure 2.**
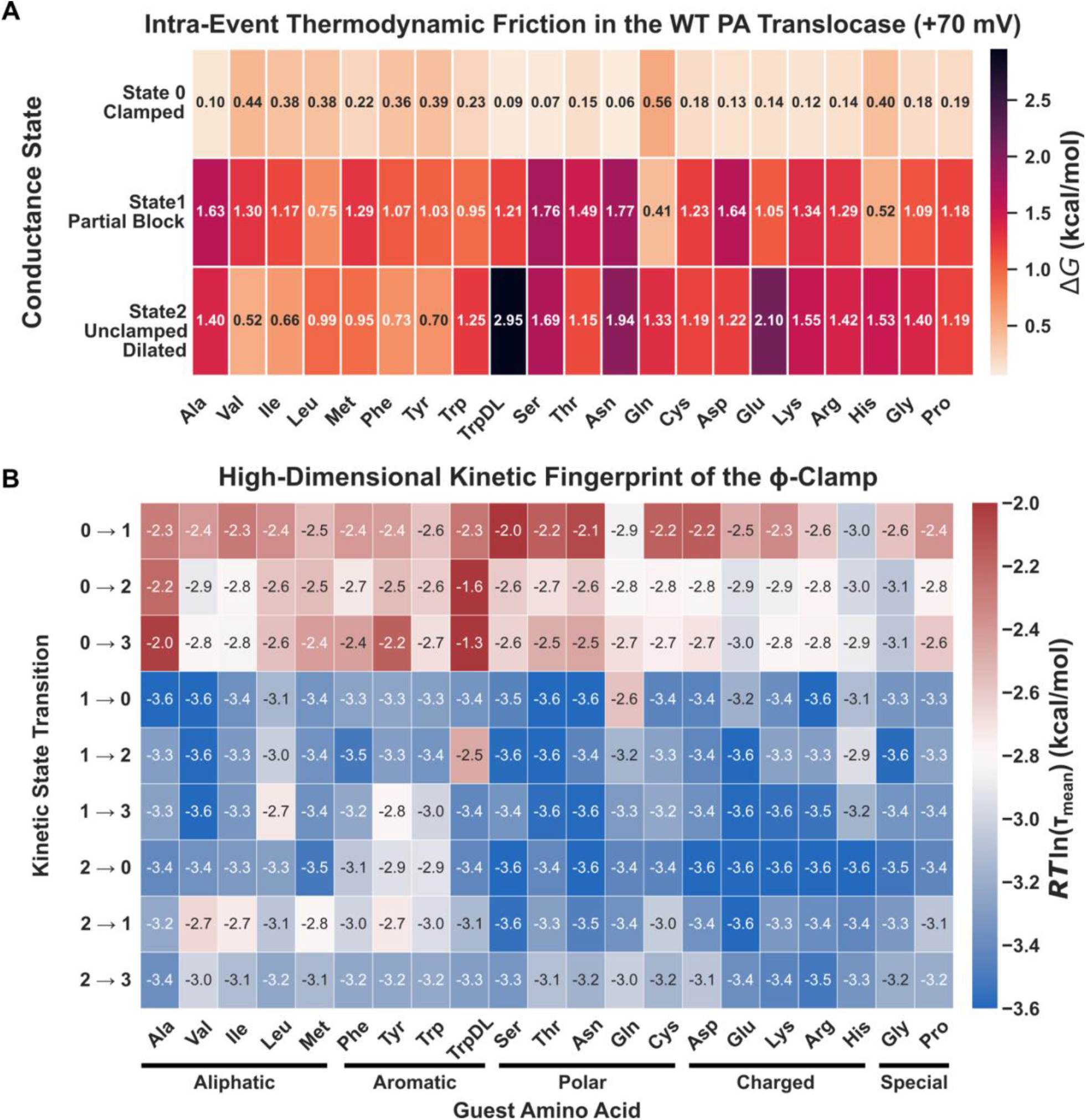
Multistate thermodynamics and kinetics of peptide translocation. **(A)** Heat maps rendering the quasi-thermodynamic state occupancies (Δ*G*) given by each indicated guest-host peptide residing in distinct conformational sub-states. **(B)** Kinetic activation energies (*RT* ln τ_mean_) required to transition between these states. In either panel, the 20 canonical amino acid guest-host peptides are shown alongside the stereochemical variant, TrpDL.

### Stereochemical Constraints and the Helix-Compression Mechanism

The stark divergence in thermodynamic and kinetic behavior between canonical l-Trp and the alternating d,l-Trp stereoisomer (TrpDL) reveals a precise structural determinant governing ϕ-clamp dilation. Quasi-thermodynamic state occupancy analysis **(Fig. 2A)** demonstrates that while l-Trp maintains a dynamic equilibrium capable of accessing the dilated sub-state (State 2, Δ*G* = 1.25 kcal/mol), TrpDL incurs a very large thermodynamic penalty (Δ*G* = 2.95 kcal/mol) against clamp dilation, locking the rigid backbone into the deep State 0 blockade.

This severe stereochemical penalty strongly supports a “helix-compression” model (*18*) for translocase ungating. To physically trigger the allosteric dilation of the F427 trapdoor to State 2, bulky analytes must transiently compress along their longitudinal axis, a structural feat natively accomplished via localized α-helix formation. While l-Trp possesses the canonical stereochemistry requisite for transient helical compression, the alternating d-stereocenters in TrpDL act as absolute helix-breakers, destroying the right-handed hydrogen-bonding network.

Consequently, ϕ-clamp ungating is deterministically governed by a dual-factor biophysical lock: it is a function of both the side-chain aromaticity (which defines the steric resting friction) and the backbone helicity (which provides the structural compression necessary to force the ϕ-clamp trapdoor open). A correlation of molecular properties of amino acids to the kinetic parameters shows a significant relationship between number of aromatic rings and the corresponding mechanism **(Supplementary Dataset 3)**. Therefore, because the TrpDL guest-host peptide possesses the extreme steric bulk of the indole rings but is physically prohibited from helical compression, it fails to ungate the translocase, resulting in the extreme State 0 → State 2 kinetic activation barriers observed in the transition matrix **(Fig. 2B)**. This stereochemical discrimination proves that the PA translocase functions as an exquisitely sensitive, multi-dimensional biophysical caliper.

### Decoding the Proteome via Physics-Informed Machine Learning (PIML)

To solve the inverse problem of proteomics – translating these kinetic fluctuations back into primary sequence identity – we deployed a PIML architecture. Rather than relying on simple amplitude matching, we engineered a hierarchical XGBoost “parallel specialist” ensemble scheme **(Supplementary Fig. S3)**. Level 1 classifiers were trained specifically to isolate orthogonal physical properties of the analytes (aromaticity, charge, hydrophobicity, and helix propensity) directly from the >60-dimension feature vector, including kinetic dwell transition matrices and other biophysical parameters describing each translocation event **(Table S1)**. These biophysical probabilities were subsequently fused by a Meta-Classifier to render the final 21-class prediction.

Tested on the full 20-amino-acid guest-host peptide library plus the stereoisomer TrpDL, with a 35 millisecond minimum-dwell time constraint to isolate deep thermodynamic friction, this parallel specialist ensemble achieved 95.81 (±0.04)% overall empirical accuracy **(Fig. 3)**. Crucially, the PIML architecture resolved stereoisomeric, isobaric, and isosteric residues that fundamentally confound traditional static nanopores and mass spectrometry. For example, the algorithm separated leucine and isoleucine (which possess identical masses and spatial volumes) by decoding the differential kinetic friction exerted by their respective branched aliphatic chains against the ϕ-clamp. This empirical benchmark demonstrates that exploiting, rather than filtering, translocase gating dynamics is the mathematical prerequisite for *de novo* peptide sequencing.

**Figure 3.**
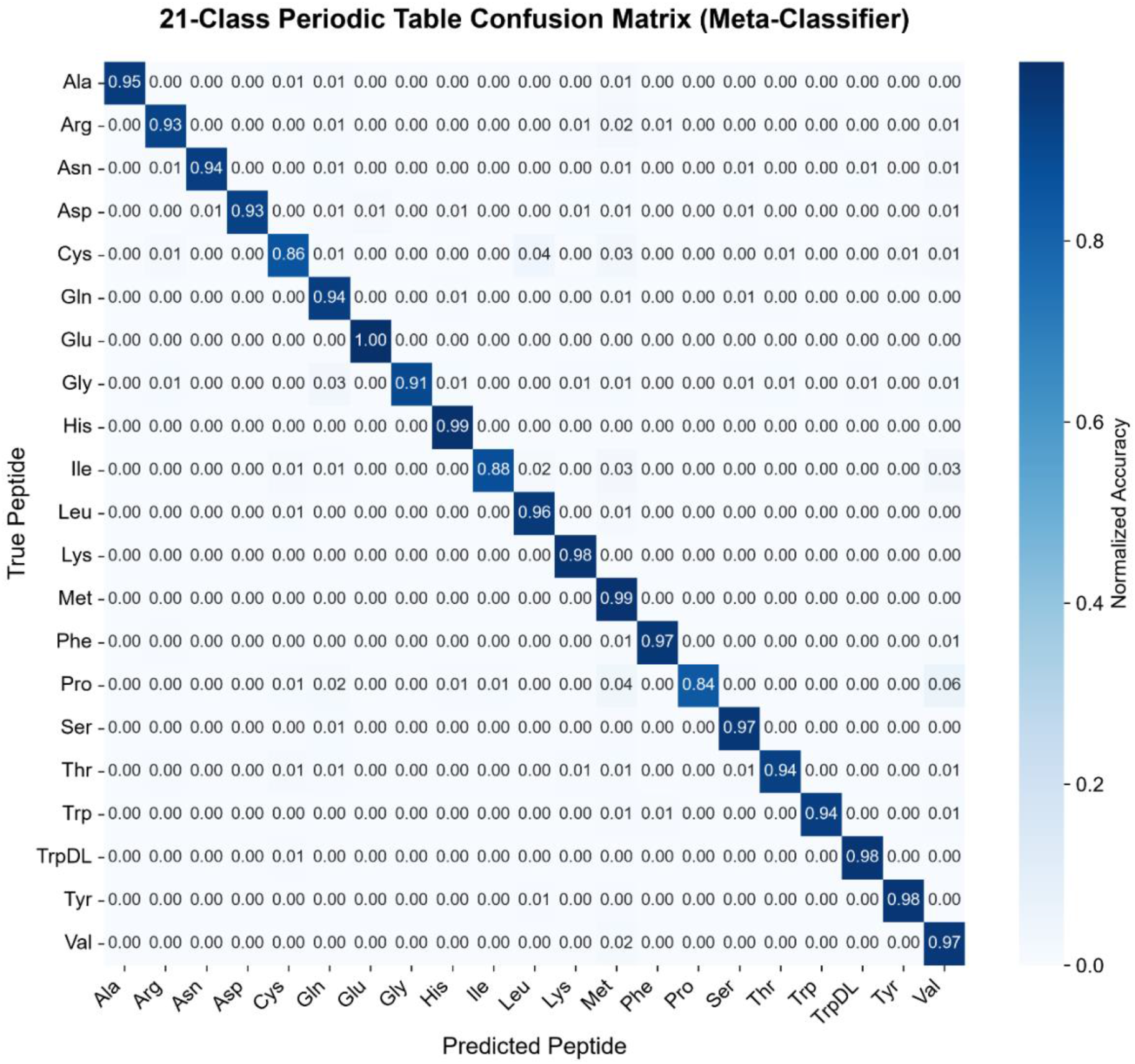
Label-free classification of the proteome with dynamical sensing and PIML. Row normalized confusion matrix for parallel specialist XGBoost model for 20 canonical amino acids as guests (X) in the peptide KKKKKXXSXX plus one stereoisomer, TrpDL. >600,000 total events were evaluated. Accuracy was 95.81 (±0.04)%, and macro-averaged F1-score was 0.9511 (±0.0005) for *N*=3 replicate train-tests of the model. Peptide concentration was 20 nM, and translocation conditions were +70 mV, 100 mM KCl, pH 5.6 symmetric.

To establish an objective benchmark for this performance, we subjected published high-bandwidth aerolysin datasets acquired from an XR_7_ guest-host peptide library (*3*) to an uncurated single-stage XGBoost pipeline **(Supplementary Fig. S4)**. Stripped of manual 2D-thresholding, the passive spatial caliper failed to resolve these isobaric overlaps, hitting a mathematical accuracy ceiling of 66.4%. For proper comparison using a simplified classifier, PA and its guest-host peptide library for the 20 canonical amino acids (K_5_XXSXX) were subjected to the same single-stage XGBoost pipeline and the resulting accuracy was 90.6 (±0.06)% **(Supplementary Fig. S5)**. This direct empirical comparison suggests that exploiting the time-domain kinetic friction of a dynamical translocase, rather than filtering it, provides a richer source of information for *de novo* peptide sequencing.

### Label-Free Resolution of Native Clinical Biomarkers

While poly-cationic guest-host systems are essential for establishing thermodynamic calibration models, true clinical proteomics requires the label-free detection of native sequences. To address and experimentally evaluates the “lysine trap” critique, the assumption that translocases require synthetic, uniform charge tails to function, we challenged the WT PA pore with a panel of untagged, native clinical biomarkers. Analytes mimicking standard “bottom-up” proteolytic fragments were driven through the pore under symmetrical buffer conditions (pH 5.0). To optimize the signal-to-noise ratio and residence time for these native analytes, the ionic strength was elevated to 500 mM KCl to boost absolute conductance, while the electrophoretic driving force was reduced to a +40 mV “kinetic brake” potential.

The dynamical translocase exhibited robust, intrinsic capture of these label-free native sequences at low nanomolar concentrations. Evaluated simultaneously in a comprehensive clinical matrix, the PIML pipeline successfully classified the oncogenic KRAS G12D neoantigen (12-mer: LVVVGADGVGKS) against the KRAS Wild-Type fragment (12-mer: LVVVGAGGVGKS), proving the system’s capacity for single-point mutation resolution in a native backbone. The platform accurately discriminated the prohormone angiotensin I (10-mer: DRVYIHPFHL) from its active cardiovascular metabolite, angiotensin II (8-mer: DRVYIHPFHL). Furthermore, the algorithm resolved the inflammatory peptide bradykinin (9-mer: RPPGFSPFR), demonstrating that the pore efficiently handles structurally rigid, proline-heavy, and highly cationic native targets. Operating on this multi-class, label-free clinical panel, the algorithm achieved a staggering 98.02 (±0.05)% overall accuracy and a 0.9771 (±0.0007) macro F1-score **(Fig. 4)**. These results support the hypothesis that the intrinsic affinity of the PA translocase’s α- and ϕ-clamps natively captures, ratchets, and decodes clinically relevant polypeptides without the need for artificial DNA handles or helicase motors.

**Figure 4.**
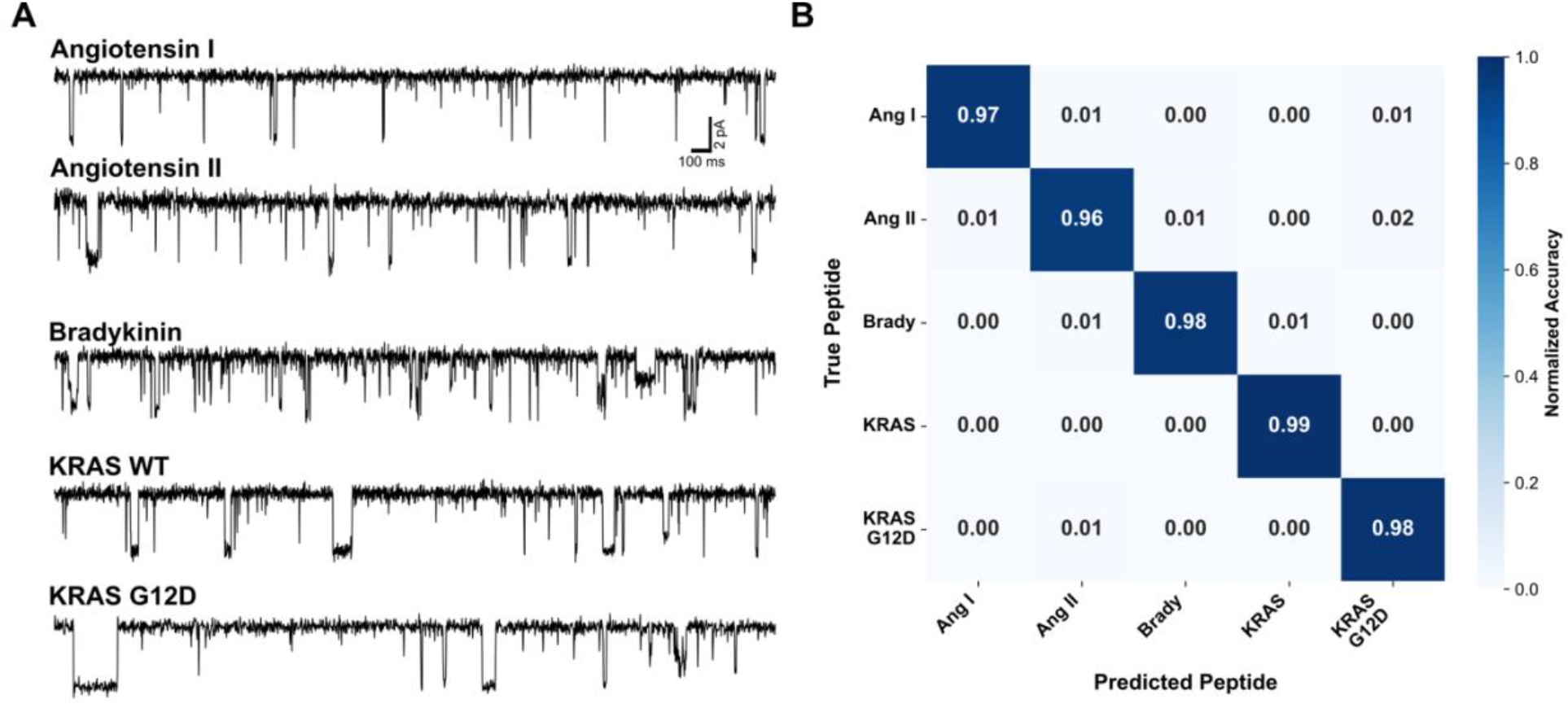
High-fidelity label-free clinical biomarker sensing. **(A)** Translocation record samples for angiotensin I, angiotensin II, bradykinin, KRAS, and KRAS G12D neoantigen. Scalebar is 2 pA by 100 ms. Peptide concentration was 20 nM, and translocation conditions were +40 mV, 500 mM KCl, pH 5.0 symmetric. **(B)** Row normalized confusion matrix for single-stage XGBoost model for 30,423 events. Accuracy was 98.02 (±0.05)%, and macro-averaged F1-score was 0.9771 (±0.0007) for *N*=3 model trainings. Three or more single pore/membrane observations were made per biomarker peptide.

## Conclusion

The realization of single-molecule proteomics has long been constrained by the fundamental limitations of static nanopores, which act as passive volumetric calipers requiring complex DNA-handle chemistries and high-micromolar analyte concentrations. Here, we establish that a dynamical translocase decisively overcomes these barriers. By actively docking and “breathing” around translocating analytes, the anthrax PA nanopore provides an exquisitely sensitive, completely label-free bio-interface operating in the low nanomolar range. This architectural shift – from a passive conduit to an active, deterministic machine – transitions single-molecule proteomics from a highly engineered laboratory phenomenon into a clinically scalable diagnostic platform.

We demonstrate that accurate sequence decoding relies on extracting the deep thermodynamic friction generated between the analyte and the ϕ-clamp, surpassing simply measuring static excluded volume. By imposing a 35-millisecond minimum-dwell time constraint, we ensure our PIML architecture captures the rich, multi-state Markovian transitions that define the physical chemistry of the peptide. Furthermore, we define the optimal substrate parameters for the dynamic sensing of peptide analytes. Unlike previous applications utilizing extended 50-residue substrates that continuously occlude the pore (*18, 27*), our system excels with short, 8-to 15-residue standard proteolytic fragments, allowing the ϕ-clamp to fully gate and generate maximum kinetic telemetry. While purely electrophoretic (Δψ) driving forces benefit from substrates possessing a nominal positive charge, this time-domain approach flawlessly resolves isobaric targets (e.g., leucine and isoleucine) that remain mathematically intractable to static nanopores and standard mass spectrometry.

The integration of a dynamical translocase with PIML represents the physical prerequisite for next-generation proteomics. Looking forward, this highly tunable platform is ideally positioned to resolve the subtle 300+ PTMs and epigenetic markers whose structural perturbations are effectively invisible to traditional analytical techniques. By deploying these dynamic reading heads onto high-throughput, multiplexed CMOS sensor arrays, future efforts will focus on isolating specific, un-tagged clinical biomarkers from dense, heterogeneous lysates – finding the diagnostic “needle in the haystack.” Ultimately, by compiling the exact thermodynamic transition matrices of the proteome, this dynamical engine establishes the foundational biophysics required to achieve true, *de novo* label-free single-molecule protein sequencing.

## Methods Summary

Single-channel electrophysiological recordings of guest-host peptide translocations through the anthrax toxin PA nanopore were acquired to map discrete, multi-state kinetic transitions. Quasi-thermodynamic and kinetic analysis were performed on extracted dwell time transition matrices for all peptides. Translocation events were processed using a custom PIML framework, extracting a >60-dimensional vector of thermodynamic and temporal features per event (e.g., dwell time transition matrices, state probabilities, etc.) to train a multi-class XGBoost classifier (*28*) **(Table S1)**. To establish an objective, static-pore benchmark, legacy high-bandwidth aerolysin datasets (*3*) were re-analyzed using an unsupervised, un-curated machine learning pipeline to determine the empirical accuracy ceiling bounded by 1D excluded volume.

Detailed protocols describing protein purification, electrophysiology, feature extraction matrices, ML hyperparameters, and bioinformatic scripts are provided in the **Supplementary Information**.

## Acknowledgments

The authors thank the department for stimulating discussions.

## Author Contributions

J.E.T. designed the experiments, collected data, analyzed results, and wrote the manuscript. P.S. contributed machine learning expertise and evaluated data analysis.

B.A.K. conceived of the PIML pipeline, designed experiments, and wrote the manuscript.

## Data, Code, and Materials Availability

Anthrax toxin raw data, custom Python classes and scripts utilized for data processing, machine learning analysis, and figure generation are publicly available via Zenodo (DOI: 10.5281/zenodo.21982624). The basic preprocessing architecture and machine learning codebase are also available on Github (https://github.com/bakrantz/Pept-Class). The Ouldali et al. primary data utilized in this study were derived from publicly available datasets (DOI: 10.13012/B2IDB-4905767_V1).

## List of Supplementary Materials

Materials and Methods

Table S1

Figs. S1 to S5

Datasets S1 to S3

## Declaration of AI and AI-Assisted Technologies in the Writing Process

During the preparation of this work, the authors used Google Gemini and Google Search to assist with literature research, refine manuscript text for clarity and readability, and optimize custom computational code. After using these tools/services, the authors reviewed and edited the content as needed and take full responsibility for the ultimate content and integrity of the publication.

